# Inorganic polyphosphate abets silencing of a sub-telomeric gene cluster in fission yeast

**DOI:** 10.1101/2022.11.28.517416

**Authors:** Ana M. Sanchez, Angad Garg, Beate Schwer, Stewart Shuman

**Affiliations:** Molecular Biology Program, Sloan-Kettering Institute, New York, NY 10065; Gerstner Sloan Kettering Graduate School of Biomedical Sciences, New York, NY 10065; Dept. of Microbiology and Immunology, Weill Cornell Medical College, New York, NY 10065

**Keywords:** *Schizosaccharomyces pombe*, polyphosphate polymerase Vtc4, transcriptional profiling

## Abstract

Inorganic polyphosphate (PolyP) is a ubiquitous polymer that plays sundry roles in cell and organismal physiology. Whereas there is evidence for polyphosphate in the cell nucleus, it is unclear whether and how physiological levels of PolyP impact transcriptional regulation in eukarya. To address this issue, we performed transcriptional profiling of fission yeast *vtc4*Δ cells, which lack the catalytic subunit of the Vtc4/Vtc2 polyphosphate polymerase complex and thus have no detectable intracellular PolyP. Deleting Vtc4 elicited the de-repression of four protein-coding genes – *SPAC186*.*04c, gdt1/SPAC186*.*05, SPAC186*.*06*, and *SPAC750*.*01* – located within a sub-telomeric region of the right arm of chromosome I that is known to be transcriptionally silenced by the TORC2 complex. These sub-telomeric genes were equally de-repressed in *vtc2*Δ cells and in cells expressing polymerase-dead Vtc4, signifying that PolyP synthesis is necessary to abet TORC2-dependent locus-specific gene silencing.

## INTRODUCTION

Inorganic polyphosphate (PolyP) is an anionic linear polymer of heterogeneous length found in taxa from all phylogenetic domains. PolyP levels and polymer chain length are determined by a dynamic balance between synthesis by PolyP kinase or PolyP polymerase enzymes and catabolism by exopolyphosphatase or endopolyphosphatase enzymes and may fluctuate in response to stress or developmental and environmental cues (1). PolyP plays diverse roles in physiology: as an energy source; a phosphate reservoir during phosphate starvation; a metal chelator; a modulator of blood clotting and fibrinolysis; a post-translation protein modification (lysine polyphosphorylation); a microbial virulence factor; a signaling molecule (1-6).

PolyP is especially abundant in yeast cells, e.g., the intracellular concentration of inorganic polyphosphate in budding yeast grown in phosphate-replete medium is 230 mM (with respect to phosphate residues) as compared to 23 mM for orthophosphate (7). Yeast PolyP is produced by a heterotrimeric membrane-associated VTC complex that synthesizes PolyP and simultaneously imports the PolyP into the yeast vacuole (8-10). Budding yeast has two VTC complexes: the VTC associated with the vacuole consists of Vtc4, Vtc3, and Vtc1 proteins; the VTC associated with the endoplasmic reticulum and nuclear envelope comprises Vtc4, Vtc2, and Vtc1 subunits (8,10). Fission yeast has a single heterotrimeric VTC complex that includes a Vtc2 subunit.

Vtc4 is the catalytic subunit of the PolyP polymerase; it consists of a cytoplasm-facing N-terminal SPX domain, a central polymerase domain, and a C-terminal membrane anchor domain (8). The SPX domain binds and senses the inositol pyrophosphate (IPP) signaling molecules IP_7_ and IP_8_ that stimulate PolyP synthesis by VTC (11-14). The Vtc4 polymerase domain, which catalyzes manganese-dependent transfer of an NTP γ-phosphate to an inorganic pyrophosphate or phosphate primer (9), is a member of the triphosphate tunnel metalloenzyme (TTM) family (15,16). Vtc2 and Vtc3 are paralogs homologous to Vtc4, but their TTM domains are catalytically inactive.

In budding yeast, vacuolar polyphosphate comprises ∼80% of the total PolyP content. There exists a pool of nuclear PolyP (dependent on Vtc4) that persists in yeast cells engineered so that the intra-vacuolar pool of PolyP is depleted (4,17). These findings raise the question of whether PolyP plays a role in nuclear transactions, especially in gene expression. To our knowledge, there is scant information on whether physiological levels of PolyP impact transcriptional regulation in eukarya. To rectify this knowledge gap, we performed transcriptional profiling of fission yeast *vtc4*Δ cells that have no detectable intracellular PolyP (13,14).

## RESULTS AND DISCUSSION

### Impact of PolyP on the fission yeast transcriptome gauged by RNA-seq

cDNAs obtained from three biological replicates, using poly(A)^+^ RNA from wild-type and *vtc4*Δ cells grown to mid-log phase in YES medium at 30°C, were sequenced. Read densities for individual genes were highly reproducible between biological replicates (Pearson coefficients of 0.98 to 0.99). A cutoff of ±2-fold change in normalized transcript read level and an adjusted p-value of <0.05 were the criteria applied to derive a list of differentially expressed annotated loci in the *vtc4Δ* mutant versus the wild-type control. We then focused on differentially expressed genes with average normalized read counts ≥100 in either strain to initially exclude transcripts that were expressed at very low levels in vegetative cells. We thereby identified 36 protein-coding genes that were down-regulated and 6 protein-coding genes that were up-regulated by these criteria in *vtc4Δ* cells (Fig. 1). All 36 genes in the former set were down-regulated modestly, between 2-fold and 4-fold *vis-à-vis* wild-type. Among the *vtc4*Δ down-regulated genes were those encoding enzymes of glycolysis and sugar metabolism: enolase (Eno102 and Eno101), glyceraldehyde-3-phosphate dehydrogenase Gpd3, glucose dehydrogenase Gcd1, transaldolase Tal1, and phosphoketolase SPBC24C6.09c. Among the up-regulated gene set, 3 coding transcripts were very strongly up-regulated: *gdt1(SPAC186*.*05c)* by 111-fold; *SPAC186*.*06* by 52-fold; and *SPAC750*.*01* by 46-fold (Fig. 1). The 3 other transcripts were increased by only 2-fold in *vtc4*Δ cells. At this stage, the RNA-seq data suggested that the presence of PolyP in wild-type cells might strongly repress the expression of a very narrow set of fission yeast transcripts. Therefore, we reset the cut-off criteria at a four-fold change in *vtc4*Δ versus wild-type and imposed no minimum threshold for read counts. This revealed one additional gene upregulated in *vtc4*Δ cells: *SPAC186*.*04c* by 46-fold (Fig. 1).

**Figure 1.**
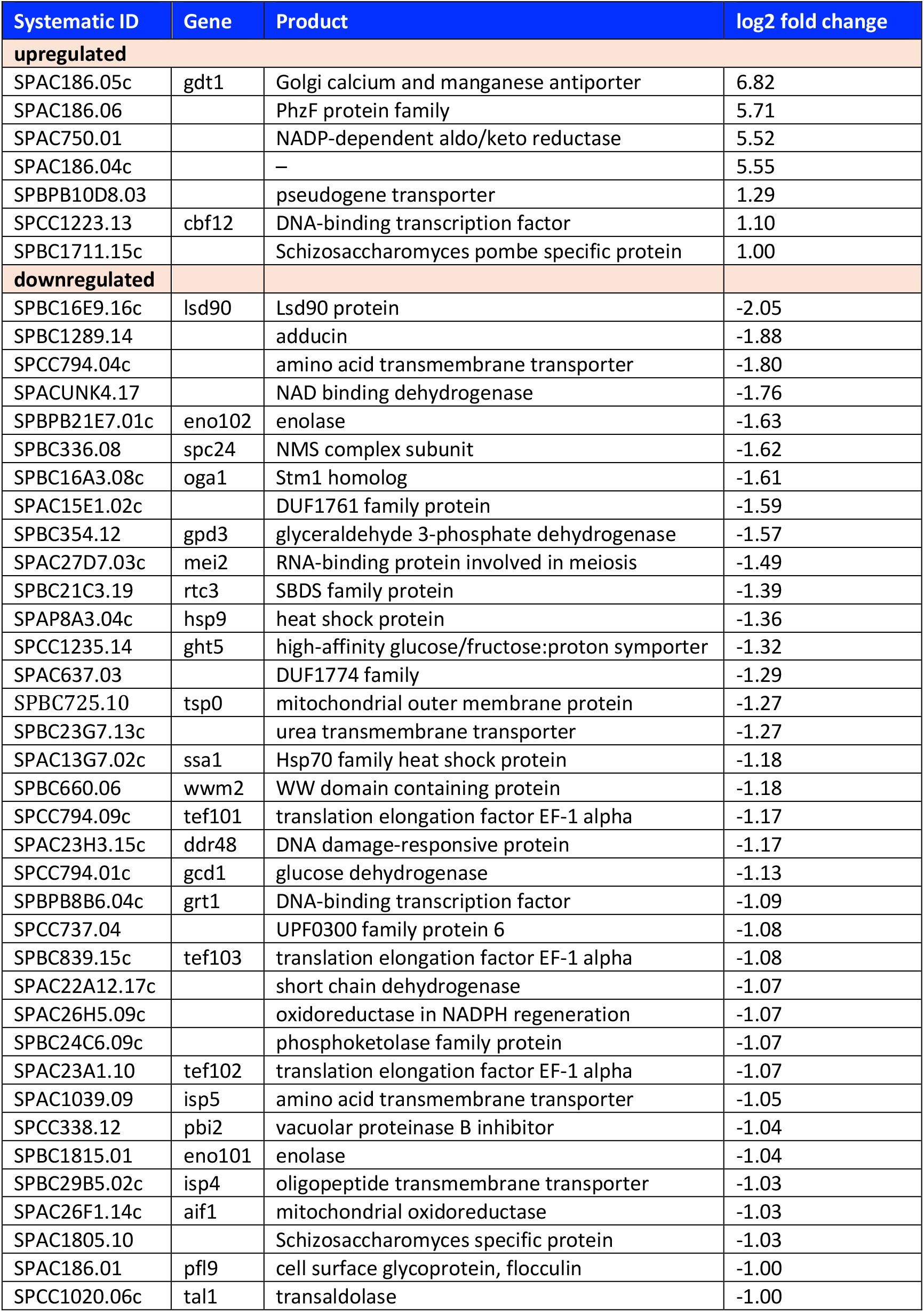
Impact of *vtc4* deletion of the fission yeast coding transcriptome. List of 7 and 36 protein-coding genes that were up-regulated and down-regulated, respectively, at least two-fold in *vtc4Δ* cells compared to the wild-type strain. The log2 fold changes are shown.

### Vtc4 repressed genes are colocalized in a sub-telomeric region

The protein products of the four *vtc4*Δ up-regulated genes appear unconnected functionally: *gdt1* encodes a Golgi calcium and manganese transporter (18); SPAC186.06 is a yeast homolog of phenazine biosynthesis enzyme PhzF (19); SPAC750.01 is a putative NADP-dependent aldo/keto reductase; SPAC186.04c is of unknown function. Their unifying property is that they are located physically within a sub-telomeric region of the right arm of chromosome I that is known to be transcriptionally silenced during unstressed vegetative growth (20). The RNA-seq read counts across the cluster of adjacent *SPAC186*.*04c, gdt1*, and *SPAC186*.*06* genes in wild-type and *vtc4*Δ cells show that the up-regulated RNA spans the entire predicted ORF and flanking UTRs, the dimensions of which can be surmised from the *vtc4*Δ RNA-seq profiles (Fig. 2).

**Figure 2.**
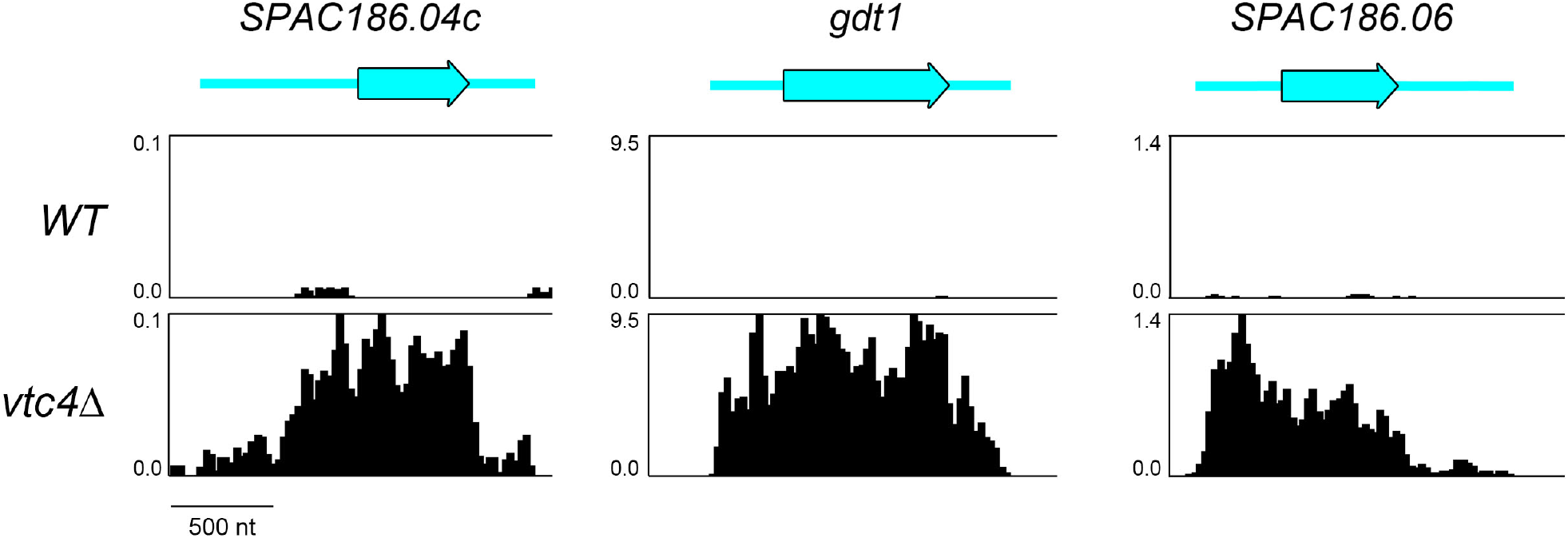
*vtc4*Δ de-represses expression of a sub-telomeric gene cluster. Strand-specific RNA-seq read densities (counts/base/million, averaged in a 25-nt window) of the indicated strains are plotted on the y-axis as a function of position across the loci (x-axis). The read densities were determined from cumulative counts of all three RNA-seq replicates for each *S. pombe* strain. The y-axis scale for each track is indicated. The common x-axis scale is shown on the bottom left. The individual ORFs are labeled and shown to scale as black-outlined thick cyan arrows. The 5’ and 3’ UTRs are depicted as thin cyan bars.

### Vtc4 repressed genes coincide with those silenced by TORC2 and Gad8

Weisman and colleagues have established that: (i) the fission yeast TORC2 complex (containing the Tor1 protein kinase) and its downstream effector protein kinase Gad8 are present in the nucleus and bound to chromatin (21,22); and (ii) TORC2 and Gad8 function to silence the expression of a set of 7 sub-telomeric genes on chromosomes I and II (20) that, not coincidentally, includes all four of the genes up-regulated by *vtc4*Δ. These sub-telomeric genes are strongly up-regulated in *tor1*Δ and *gad8*Δ cells; they are also de-repressed in *ste20*Δ and *ryh1*Δ cells that respectively lack the TORC2 subunit Ste20 and the Ryh1 GTPase that activates TORC (20). The de-repression of the sub-telomeric loci in *tor1*Δ and *gad8*Δ cells correlates with: (i) loss of the repressive H3K9Me2 chromatin mark over the loci; (ii) gain of the H3K4me3 and H4K16Ac activation marks; and (iii) increased locus occupancy by Pol2 (20). Further studies showed that increased expression of the *gdt1(SPAC186*.*05c), SPAC186*.*04c*, and *SPAC186*.*06* genes in *tor1*Δ cells depends on Pol2 transcription activators Leo1, Med1, and Gcn5 (23). Our transcriptional profiling implicates the VTC complex as a collaborator in TORC2 silencing of the sub-telomeric four-gene cluster on chromosome I.

#### VTC subunits and PolyP polymerase activity are required for localized gene silencing

A salient question is whether the transcriptional impact of *vtc4*Δ on sub-telomeric gene silencing is a consequence of the absence of PolyP or the interdiction of a hypothetical function of Vtc4 other than PolyP synthesis. If the VTC complex is necessary for silencing, then we would expect that deletion of the Vtc2 subunit would phenocopy *vtc4*Δ. The requirement for catalysis by Vtc4 can be interrogated via a polymerase active site mutation R262A-R264A in Vtc4 that eliminates PolyP in fission yeast (14). To gauge expression of the four sub-telomeric genes in wild-type, *vtc4*Δ, *vtc2*Δ, and *vtc4-(R262A-R264A)* genetic backgrounds, we performed transcript-specific RT-qPCR of total RNA isolated from these four strains. RT-qPCR analysis of *act1* mRNA was included as a control. The sub-telomeric transcript levels were normalized to *act1* for each sample and are plotted in Fig. 3, such that the changes in transcript levels in the *vtc* mutants are normalized to transcript levels of wild-type cells. The findings were as follows: (i) RT-qPCR affirmed that *gdt1, SPAC186*.*04c, SPAC186*.*06*, and *SPAC750*.*01* were strongly de-repressed in *vtc4*Δ cells; (ii) similar extents of derepression were observed in *vtc2*Δ and *vtc4-(R262A-R264A)* cells, signifying that PolyP synthesis by the VTC complex is required for silencing of these sub-telomeric genes.

**Figure 3.**
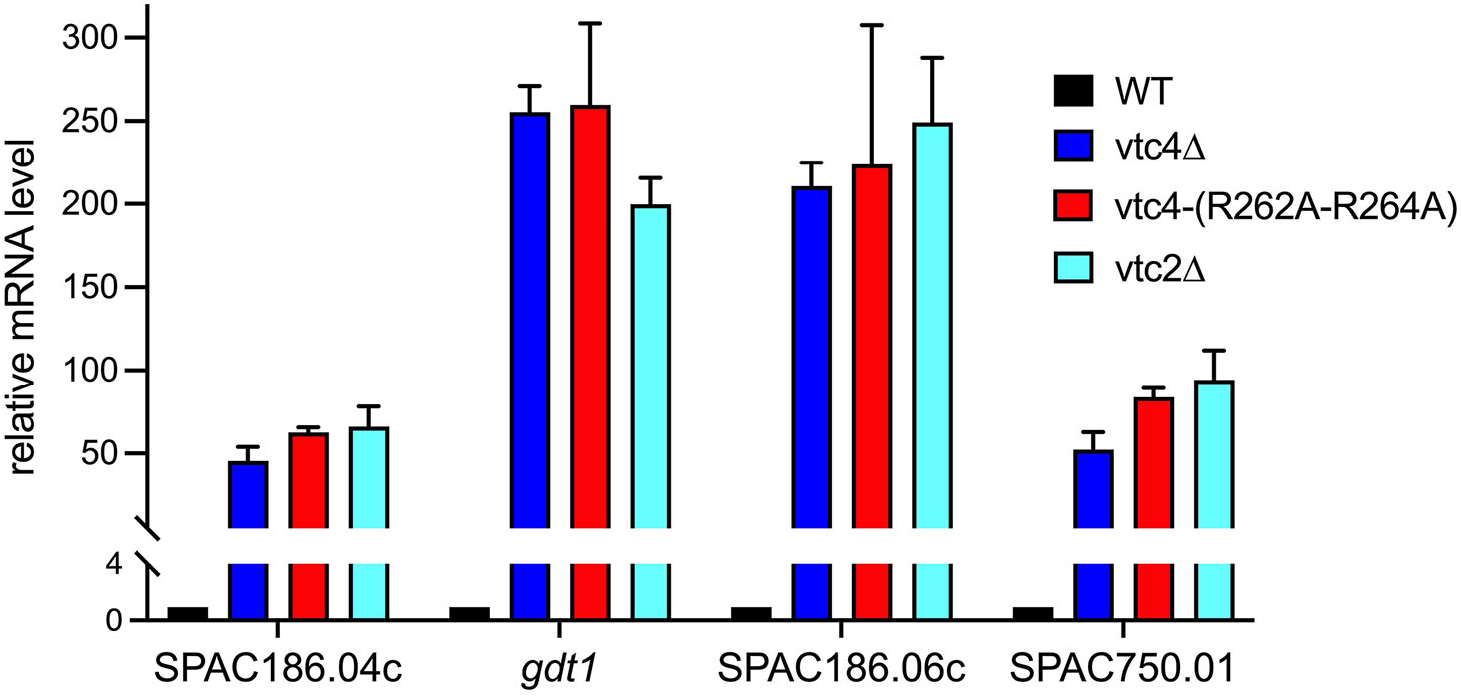
VTC PolyP polymerase activity is required for silencing the sub-telomeric genes. The analysis of mRNA levels for *SPAC186*.*04c, gdt1, SPAC186*.*06c, SPAC750*.*01* in *WT, vtc4Δ, vtc4-(R262A-R264A)*, and *vtc2Δ* yeast strains was performed as described in Methods. The level of each transcript was normalized to that of *act1* for the same RNA sample. The bar graph shows relative change in mRNA levels in *vtc4Δ, vtc4-(R262A-R264A)*, and *vtc2Δ* relative to the wild-type control (defined as 1.0). Each datum in the graph is the average values of RT-qPCR analyses of RNA from three independent yeast cultures; the error bars denote SEM.

#### Role of IPP-binding to Vtc4 and Vtc2 SPX domains in PolyP synthesis

PolyP synthesis by the budding yeast VTC complex in isolated vacuoles is stimulated ∼20-fold by sub-micromolar concentrations of IPPs, with 1,5-IP_8_ being at least 20-fold more potent than 5-IP_7_ or 1-IP_7_ based on EC_50_ (12). In fission yeast, cellular PolyP levels correlate with the levels of IP_8_ synthesized by IPP kinase Asp1, i.e., the PolyP content of *asp1*Δ and *asp1-D333A* (IPP kinase-inactive) cells, which fail to synthesize IP_8_, is reduced compared to wild-type, albeit not depleted completely as in *vtc4*Δ (13,14). A previous study documenting the importance of IPP binding to the VTC complex for PolyP synthesis by isolated *S. cerevisiae* vacuoles was based on the effects of simultaneously mutating the IPP-binding residues of both SPX domains of the VTC complex, which abolished IP_7_-stimulated PolyP synthesis (11). To query the role of the individual Vtc4 and Vtc2 SPX domains, we exploited fission yeast mutants with triple-alanine substitutions, Y22A-K26A-K30A, in the IPP binding sites of Vtc4 and Vtc2, respectively. We then proceeded to generate a *vtc4-(Y22A-K26A-K30A) vtc2-(Y22A-K26A-K30A)* strain in which both IPP-binding sites of the fission yeast VTC complex were mutated. We employed an optimized protocol to recover polyphosphates from whole-cell extracts (24), which were analyzed by polyacrylamide gel electrophoresis and stained with toluidine blue to compare the PolyP content of wild-type cells to that of cells with Vtc4 and Vtc2 SPX domain mutants. *vtc4*Δ and *asp1*Δ extracts were analyzed in parallel as controls. The pertinent findings were as follows: (i) PolyP content of the *vtc4-(Y22A-K26A-K30A)* and *vtc2-(Y22A-K26A-K30A)* strains was similar to wild-type cells; (ii) PolyP content of the *vtc4-(Y22A-K26A-K30A) vtc2-(Y22A-K26A-K30A)* strain was diminished, especially the longer population of PolyP chains; and (iii) the impact of simultaneously mutating both VTC SPX domains was similar to eliminating IP_8_ synthesis in the *asp1*Δ strain (Fig. 4). We infer that IP_8_ binding to either the Vtc4 or Vtc2 SPX domain suffices to elicit IPP-stimulation of PolyP synthesis.

**Figure 4.**
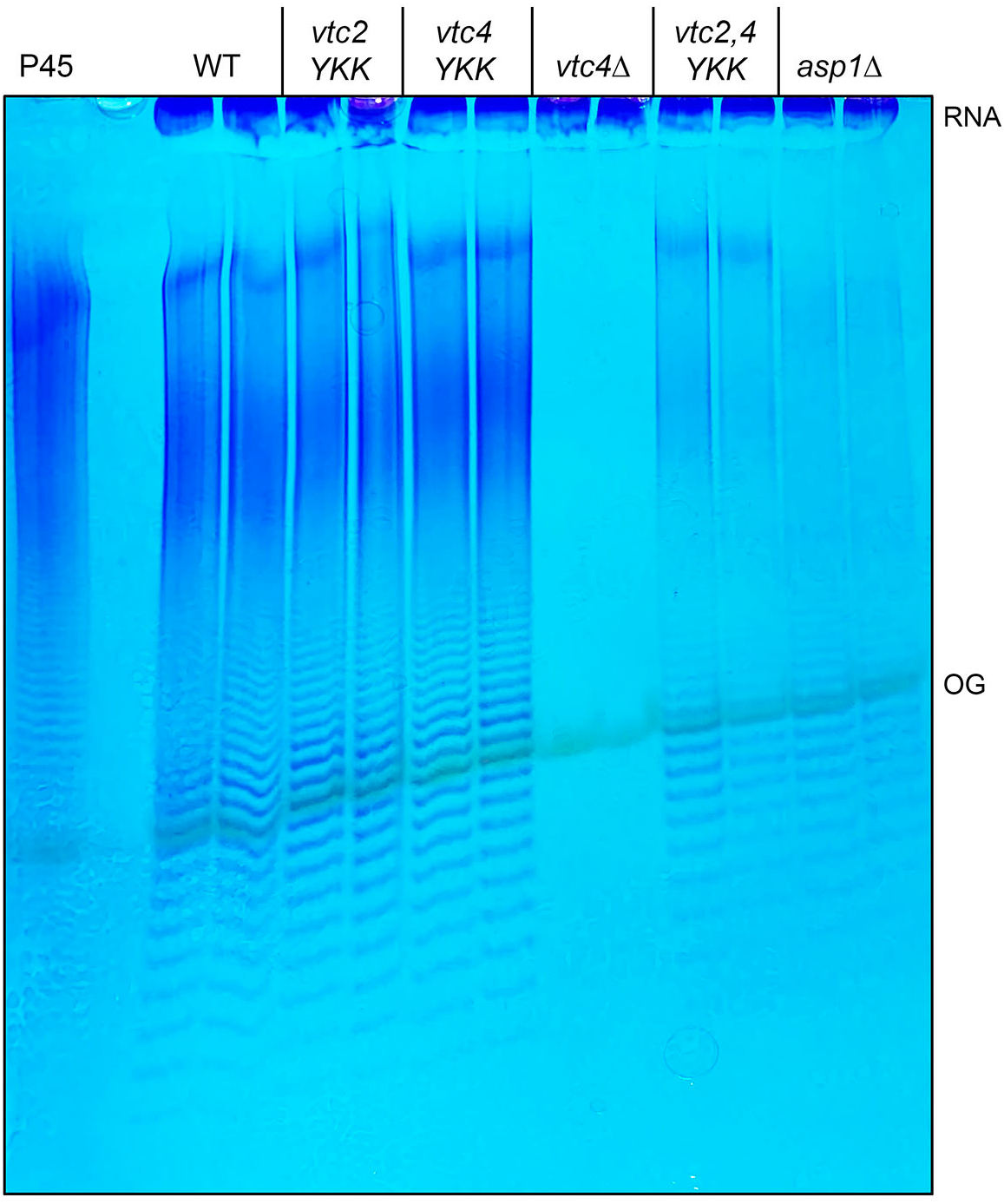
Effect of VTC SPX domain IPP binding site mutations on cellular PolyP. Total polyphosphate prepared from the indicated fission yeast strains was analyzed by PAGE and visualized by staining with toluidine blue. Lane P45 contains commercial polyphosphate (average chain length 45; Sigma S4379). The stained material near the top of the gel is RNA. The position of the Orange G dye marker (OG) is indicated.

#### An emerging appreciation of PolyP as a locus-specific gene silencer

The results presented here anent the de-repression of a cluster of normally silenced subtelomeric genes in fission yeast when PolyP synthesis is interdicted genetically resonate with the recent discovery by Jakob and colleagues that deletion of PolyP kinase in *E. coli* elicits up-regulation of transcription of normally silenced chromosomal regions containing mobile genetic elements and prophages (25). Their experiments indicate that PolyP collaborates with bacterial Hfq and AT-rich DNA to nucleate phase-separated condensates. They propose that PolyP drives the formation of localized heterochromatin in bacteria (25).

Analogous genetic studies of PolyP physiology in mammalian cells are not feasible at present, because the mammalian enzyme(s) responsible for PolyP synthesis are not known (2). A surrogate approach has been to overproduce PolyP in mammalian cells via ectopic expression of bacterial PolyP kinase and document the effects thereof. When this maneuver was applied to human cells, it was found that: (i) high levels of long-chain PolyP accumulated in multiple intracellular compartments; and (ii) 313 genes were down-regulated and 47 genes were upregulated in PolyP kinase-expressing cells, by the criterion of a statistically significant 25% difference versus non-expressing cells (26). If a ≥2-fold difference cut-off is applied, then 102 genes were down-regulated and 8 genes were upregulated. There are caveats to this approach when it comes to inferences about PolyP function. To wit: (i) the levels of PolyP achieved are excessive, hence non-physiological; (ii) the PolyP may localize to intracellular sites where it is not normally present; and (iii) the PolyP that does accumulate is skewed toward very long chains. Indeed, studies in budding yeast indicate that forced accumulation of non-physiological levels of polyphosphate outside the vacuole and membrane compartments, achieved via expression of a bacterial PolyP kinase, is *per se* cytotoxic (10) Coupling of VTC-mediated polyphosphate polymerase activity to vacuolar or intramembrane import of the polyphosphate product is a means to avoid such toxicity (10).

#### How might PolyP locally impact fission yeast transcription regulation

The present data instate a role for fission yeast PolyP in localized gene silencing, presumably as a participant in the TORC2 pathway of sub-telomeric silencing discovered by the Weisman lab. Based on available knowledge, we can speculate on at least two ways in which PolyP may accomplish this. First, PolyP can exert effects on cell physiology via non-enzymatic lysine polyphosphorylation of target proteins in vivo, including nuclear proteins such as DNA topoisomerase I, Nsr1, and ribosome biogenesis factors (27-29). Indeed, among the validated targets of in vivo lysine polyphosphorylation in budding yeast are several proteins involved in chromatin biology: histone H2AZ chaperone Chz1, histone acetyltransferase complex subunit Eaf7, nucleosome assembly factor Hpc2 (28). If lysine polyphosphorylation of nuclear proteins that establish or maintain silenced chromatin over the fission yeast chromosome I sub-telomeric cluster is important for their activity, then ablation of PolyP synthesis would elicit the observed de-repression. Given that the TORC2 pathway is necessary for silencing the cluster in fission yeast, it is possible TORC2 or pathway components acting downstream are subject to lysine polyphosphorylation. Second, taking a cue from the recent studies in *E. coli* (25), PolyP might promote the assembly of repressive factors at the affected loci, by providing a scaffold for their recruitment to form a higher order PolyP–protein– DNA assembly, with or without accompanying phase separation. Other models or mechanisms for PolyP-mediated silencing are in no way off the table. In conclusion, the initial findings here provide an impetus for further interrogation of PolyP function in fungal gene expression.

## METHODS

### Transcriptome profiling by RNA-seq

RNA was isolated from *S. pombe* wild-type and *vtc4*Δ cells that were grown in liquid YES medium at 30°C to an *A*_600_ of 0.5 to 0.6. Cells were harvested by centrifugation and total RNA was extracted via the hot phenol method. The integrity of total RNA was gauged with an Agilent Technologies 2100 Bioanalyzer. The Illumina TruSeq stranded mRNA sample preparation kit was used to purify poly(A)^+^ RNA from 500 ng of total RNA and to carry out the subsequent steps of poly(A)^+^ RNA fragmentation, strand-specific cDNA synthesis, indexing, and amplification. Indexed libraries were normalized and pooled for paired-end sequencing performed by using an Illumina NovaSeq 6000-S1 flow cell. FASTQ files bearing paired-end reads of length 51 bases (total paired reads of 19.1 million to 27.5 million per biological replicate) were mapped to the *S. pombe* genome (Pombase) using HISAT2-2.1.0 with default parameters (30). Mapped reads comprised 93% to 96% of the total reads per replicate. The resulting SAM files were converted to BAM files using Samtools (31). Count files for individual replicates were generated with HTSeq-0.10.0 [51] using exon annotations from Pombase (GFF annotations, genome-version ASM294v2; source ‘ensembl’). RPKM analysis and calculations of pairwise correlations (Pearson coefficients of 0.978 to 0.987) were performed as described previously (32). Differential gene expression and fold change analysis was performed in DESeq2 (33). Cut-off for further evaluation was set for genes that had an adjusted p-value (Benjamini-Hochberg corrected) of ≤0.05 and were up or down by at least two-fold in *vtc4*Δ versus wild-type. Genes were further filtered on the following criteria: (i) ≥2-fold up and the average normalized read count for the mutant strain was ≥100; and (ii) ≥2-fold down and the average normalized read count for the wild-type strain was ≥100. Alternatively, a cut-off of at least a four-fold up or down in *vtc4*Δ versus wild-type was set without regard to the normalized read count values, which flagged *SPAC186*.*04c* as upregulated in *vtc4*Δ cells.

### Reverse transcriptase quantitative PCR analysis

Total RNA was prepared from exponentially growing cells (three independent cultures for each yeast strain analyzed) via the hot phenol method. The RNAs were treated with DNase I, extracted serially with phenol:chloroform and chloroform, and then precipitated with ethanol. The RNAs were resuspended in 10 mM Tris HCl (pH 6.8) and 1 mM EDTA and adjusted to a concentration of 600 ng/μl. Reverse transcription was performed with 2 μg of this RNA template plus oligo(dT)_18_ and random hexamer primers by using the Maxima First Strand cDNA synthesis kit (Thermo Scientific). After cDNA synthesis for 30 min at 55°C, the reverse transcription reaction mixtures were diluted 10-fold with water. Aliquots (2 μl) were used as templates for gene-specific quantitative PCR (qPCR) reactions directed by the sense and antisense primers listed in Table S1. The qPCR reactions were constituted with the Maxima SYBR Green/ROX master mix (Thermo Scientific) and monitored with an Applied Biosystems QuantStudio 6 Flex Real-Time PCR system. The qPCR reactions were performed in triplicate for each cDNA population. The level of individual cDNAs was calculated relative to that of *act1* cDNA by the comparative Ct method (34). The actin-normalized levels of the four sub-telomeric transcripts in wild-type cells were assigned a value of 1.0 and the mRNA levels in the three *vtc* mutant were then normalized to the wild-type control value.

### PAGE assay of intracellular polyphosphate content

*S. pombe* cells were grown in YES medium at 30°C. Aliquots (5 *A*_600_ units) of logarithmically growing duplicate cultures of each strain were harvested by centrifugation, washed with cold water and stored at -80°C. The cell pellets were resuspended in 400 μl AE buffer (50 mM sodium acetate pH 5.2, 10 mM EDTA) and added to 300 μl phenol (equilibrated in 10 mM Tris-HCl, pH 8.0) plus 40 μl of 10% (w/v) SDS. The samples were mixed vigorously and incubated at 65°C for 5 min and then placed on ice. Chloroform (300 μl) was added, and the mixed samples were centrifuged for 5 min at room temperature at 17,000g. The aqueous phase (∼450 μl) was collected and extracted with phenol/chloroform and chloroform and then ethanol precipitated at -20°C overnight. The precipitates were washed with 70% ethanol, dried and resuspended in 75 μl TE (10 mM Tris-HCl, pH 7.0, 1 mM EDTA; 15 μl per 1 *A*_600_ unit of cells). Aliquots (10 μl) were supplemented with 3x Orange G dye (10 mM Tris-HCl, pH 7.0, 1 mM EDTA, 30% glycerol, 0.1% [w/v] Orange G) and then analyzed by electrophoresis through a 36% polyacrylamide gel in TBE (80 mM Tris-borate, 1 mM EDTA) at 4°C for 17 h at 5 mA. The gel was stained with toluidine blue (0.05% [w/v] toluidine blue in 20% methanol, 2% glycerol) and scanned after de-staining.

## Data Deposition

The RNA-seq data in this publication have been deposited in NCBI’s Gene Expression Omnibus and are accessible through GEO Series accession number GSE213524.

## Supplementary information

## FIGURE LEGENDS

**Table S1.**
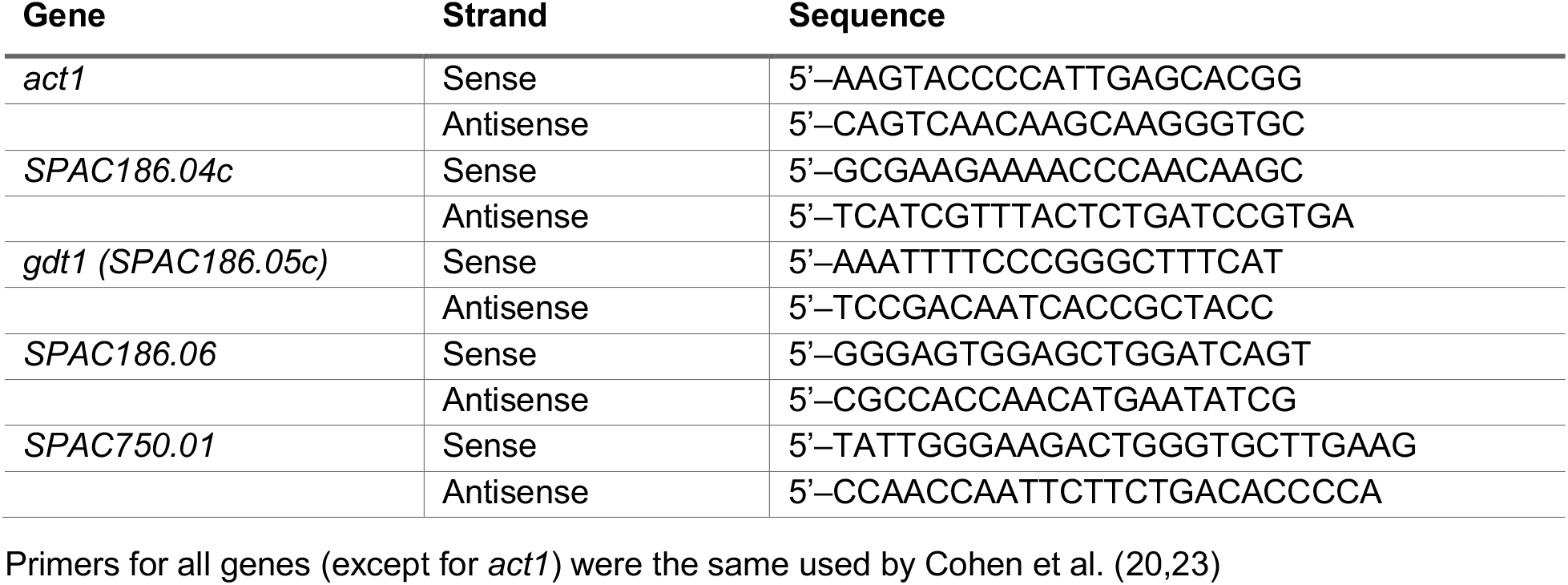
Oligonucleotide primers used for qPCR analyses

**Table S2.** Fission yeast strains used in this study

| Strain | Genotype |
| --- | --- |
| BS78 | *h+ vtc2Δ::kanMX leu1-32* |
| BS128 | *h- vtc4Δ::kanMX leu1-32* |
| BS620 | *h- vtc4-(Y22A-K26A-K30A)::kanMX leu1-32* |
| BS623 | *h+ vtc4-(R262A-R264A)::kanMX leu1-32* |
| AS1887 | *h- asp1Δ::kanMX leu1-32* |
| BS793 | *h- leu1<sup>+</sup>::vtc2-(Y22A-K26A-K30A) vtc2Δ::hygMX* |
| BS803 | *h- leu1<sup>+</sup>::vtc2-(Y22A-K26A-K30A) vtc2Δ::hygMX vtc4-(Y22A-K26A-K30A)::kanMX* |

All strains are *leu1-32 ura4-D18 his3-D1* and either *ade6-m216* or *ade6-m210*.

